# On the equivalence between agent-based and continuum models for cell population modeling. Application to glioblastoma evolution in microfluidic devices

**DOI:** 10.1101/2024.09.05.611243

**Authors:** Raquel B. Arroyo-Vázquez, Marina Pérez-Aliacar, Jacobo Ayensa-Jiménez, Manuel Doblaré

**Affiliations:** Aragón Institute of Engineering Research (I3A), University of Zaragoza; Mechanical Engineering Department Aragón Institute of Engineering Research (I3A), University of Zaragoza; Aragón Institute of Engineering Research (I3A), Institute for Health Research Aragón (IISAragón),University of Zaragoza; Aragón Institute of Engineering Research (I3A), Institute for Health Research Aragón (IISAragón), University of Zaragoza, CIBER-BBN, ISCIII, Nanjing Tech University

**Keywords:** Agent-Based Model, Continuum models, Glioblastoma, PhysiCell, Microfluidics, Mathematical modelling

## Abstract

Mathematical models are invaluable tools for understanding the mechanisms and interactions that control the behavior of complex systems. Modeling a problem as cancer evolution includes many coupled phenomena being therefore impossible to obtain sufficient experimental results to fully evaluate all possible conditions. In this work, we focus on Agent-Based Models (ABMs), as these models allow to obtain more complete and interpretable information at the individual level than other types of *in silico* models. However, ABMs, need many parameters, requiring more information at the cellular and environmental levels to be calibrated. To overcome this problem we propose a complementary approach to traditional calibration methods. We used existent continuum models able to reproduce experimental data, validated and with fitted parameters, to establish relationships between parameters of both, continuum and agent-based models, to simplify and improve the process of adjusting the parameters of the ABM. With this approach, it is possible to bridge the gap between both kinds of models, allowing to work with them simultaneously and take advantage of the benefits of each of them.

To illustrate this methodology, the evolution of glioblastoma (GB) is modeled as an example of application. The resulting ABM obtains very similar results to those previously obtained with the continuum model, replicating the main histopathological features (the formation of necrotic cores and pseudopalisades) appearing in several different in vitro experiments in microfluidic devices, as we previously obtained with continuum models. However, ABMs have additional advantages: since they also incorporates the inherent random effects present in Biology, providing a more natural explanation and a deeper understanding of biological processes. Moreover, additional relevant phenomena can be easily incorporated, such as the mechanical interaction between cells or with the environment, angiogenic processes and cell concentrations far from the continuum requirement as happens, for intance, with immune cells.

## 1 Introduction

Cancer is one of the deadliest diseases in the world. In 2022, according to the World Health Organization (WHO), approximately 20 million new cases were diagnosed, and almost 50% died from this disease. Despite all the research being carried out in this regard, it has not been possible to reduce the number of cases or the mortality associated with them. In fact, by 2050, the WHO predicts an increase in both incidence and mortality [WHO, 2024].

When studying and modeling cancer and its progression, various approaches are employed, including *in vivo, in vitro*, and *in silico* models. Each of these methods has its own set of advantages and disadvantages, so several different approaches are typically used in a combined way.

*In vivo* models, the most complex ones, aim to establish similarities with tumors and their behavior in real patients. However, they present challenges, as it is impossible to isolate and control every phenomenon present in experiments [Neufeld et al., 2021], having therefore a poor reproducibility. By contrast, *in vitro* models address some of these challenges, as they allow for the reproduction of specific biological phenomena with greater control over the experiment variables, having a higher reproducibility and leading to a deeper understanding of the underlying mechanisms. In particular, in recent years, microfluidic technology has emerged as a powerful tool for get richer and more accurate *in vitro* models [Mehling and Tay, 2014, Wu et al., 2010, Mehta et al., 2022].

Microfluidics is a technique that combines the manipulation of fluids in a network of micrometer-scale channels with 3D cell culture chambers. This technique allows to reproduce the different gradients of the tumor microenvironment and study their effect in heterogeneous cultures. All this inside small chips, with the advantages of the low cost of agents and material required, and the high resolution and sensitivity that can be achieved [Whitesides, 2006, Logun et al., 2018]. This technology gives rise to the so-called *Organ-on-chip* systems [Ma et al., 2021], which, unlike traditional Petri dishes, makes it possible to reproduce complete biological systems, with the long-term aim of replacing preclinical animal models [Low et al., 2020].

Despite all the above-mentioned advantages, *in vitro* models still have limitations when trying to dissociate intrinsically coupled phenomena, quantifying the effect of the control and internal variables involved in the experiments, testing new hypotheses or predicting the behavior of cells in “unseen” situations. For this reason, they have been progressively combined with *in silico* models (physical-mathematical models). These latter are highly effective and are currently capable of describing and simulating complex biological problems [Zhang et al., 2011, Macklin et al., 2012, Alfonso et al., 2017, Gong et al., 2017, Miranda et al., 2018, Senders et al., 2020, Randles et al., 2021], particularly when combined with high throughput screening platforms, such as microfluidic devices [Martínez-González et al., 2012, Du et al., 2016, Ayuso et al., 2017, Ayensa-Jiménez et al., 2020, Ayensa-Jiménez et al., 2022].

There are multiple types of mathematical models, but we focus here on Agent-Based Models (ABMs)[An et al., 2009]. ABMs model the coupled behavior of a set of entities, called agents, that take decisions autonomously. These decisions are regulated by a set of rules that the user defines and determine the actions of the agents for each set of external signals that they receive. For instance, in the case of tumor progression, the agents are tumor cells that are subjected to different stimuli: pressure, oxygen gradients, biochemical signals secreted by other cells, etc. Subsequently, through the set of rules introduced that define the agent model, this external information is translated into cell actions such as: migrating, modifying their volume, replicating, dying or changing their phenotype, among others.

In general, ABMs offer several advantages. They work at the individual scale allowing to analyze how local interactions give rise to global behaviors, thus capturing emergent phenomena at the population level [Yu and Bagheri, 2020]. When modeling individual cells, they provide a natural description of the system, with actual stimuli and responses observed in Biology, and with internal behavior as complex as required being easier to interpret [Wang et al., 2015]. Moreover, the ABMs inherently incorporate stochastic phenomena, allowing the uncertainty of predictions to be controlled and analyze. Finally, they have a modular structure and are more flexible, allowing to easily introduce new agents or change rules, without having to change the entire model, as well as to analyze the effect of global parameters whose biological interpretability is obscure.

ABMs have been used to study the interactions between cells and their microenvironment [Bonabeau, 2002, An et al., 2009]. In particular, when simulating tumor heterogeneity [Mirams et al., 2013, Bull et al., 2020], they allow to incorporate the interaction between cancer cells and the immune system [Gong et al., 2017, Kather et al., 2017, Ghaffarizadeh et al., 2018, Metzcar et al., 2019], or angiogenic processes [Bauer et al., 2007, Phillips et al., 2020], among other important phenomena in cancer evolution.

However, ABMs have a high computational cost, tend to be more complex, and are more dependent on their implementation [Bonabeau, 2002]. Even more dramatic, these models have many parameters, which in many cases translates into a lack of parameter identifiability [Robin et al., 2024] and the fitting task is tremendously complicated [McCulloch et al., 2022]. To solve this problem, researchers use different methodologies, such as machine learning [Cess and Finley, 2023, Lamperti et al., 2018], Gaussian processess [Dancik et al., 2010] or Bayesian optimization [Movilla et al., 2022], although parameter identification is an intrinsic challenge in ABMs.

In this work, we propose a different approach based on knowledge inheritance from previous models: We decided to use continuum models able to reproduce experimental data, sufficiently validated and with fitted parameters, to establish relationships between the parameters of the continuum and agent-based models trying to simplify the parameter fitting process for ABMs.

The derivation of equivalences between individual or agent-based and continuum models and their corresponding parameters has been a topic of extensive research in the last few years. Examples include the derivation of continuum equations for cell movement, including diffusion (random migration) from random-walk models Baker et al. [2009], Gerlee and Nelander [2012], Morselli et al. [2023] and/or chemotaxis Baker et al. [2009]. Some works, such as Deroulers et al. [2009], Murray et al. [2009] include interactions between cells, arriving to non-linear diffusion models. Other authors explored the relationship of ABMs with continuum models when the growth and migration rates are pressure-based Chaplain et al. [2020], Byrne and Drasdo [2009], Lötstedt [2021], Morselli et al. [2023]. Very recently, Stace et al. [2020], Macfarlane et al. [2022], Ardaševa et al. [2020] derived continuum models from ABMs including the dependence of migration and proliferation on the cell phenotypic state. Regarding the methodology, most authors use coarse-grain methods and derive the continuum limit of the ABM Macfarlane et al. [2022], Ardaševa et al. [2020], Gerlee and Nelander [2012], Murray et al. [2009]. Data science methods, and in particular equation learning based on sparse regression have also been used to infer continuum models from ABMs Nardini et al. [2021]. In the present study, we derive a new set of relationship between the parameters of a class of ABMs, incorporating cellular proliferation and death, as well as random and directed migration, with a class of continuum models based on reaction diffusion equations.

To illustrate this methodology, the evolution of *in vitro* glioblastoma (GB) cells is modeled. GB is a type of brain cancer that affects glial cells. According to WHO, GB is of grade IV, in its most advanced stage. It is also the most commonly occurring malignant brain tumor, with an incidence rate of 3-4 cases per 100,000 individuals per year [Louis et al., 2021, Grochans et al., 2022]. GB is highly aggressive and invasive, with an average survival rate of only 15 months following diagnosis [Lah et al., 2020, Grochans et al., 2022]. Its lethality is largely attributed to its high resistance to radiotherapeutic and chemotherapeutic treatment, as well as the inability to completely eliminate the tumoral tissue by brain surgery [Lah et al., 2020].

The evolution of this tumor is characterized by a high cell proliferation around blood vessels [Rosińska and Gavard, 2021], leading to their occlusion and the subsequent emergence of hypoxic zones [Grimes et al., 2020]. These areas may eventually develop into necrotic regions (with cells dying due to oxygen deprivation), and cells migrating towards more oxygenated ones, giving rise to travelling waves of cells known as pseudopalisades [Rong et al., 2006]. That’s why one crucial factor in the evolution of this tumor is the availability of oxygen, as it regulates tumor growth, invasiveness, and necrosis, in addition to other fundamental aspects, such as cellular resistance and adaptation [Brat and Van Meir, 2004].

To reproduce GB evolution, several models in the literature have been proposed [Swanson et al., 2007, Martínez-González et al., 2012, Ayuso et al., 2017, Ayensa-Jiménez et al., 2020]. We have chosen the continuum model of Ayensa-Jiménez et al. [2020], since it is adapted to experiments coming from microfluidic devices and, due to its flexibility and the fact that it combines random and directed migration when considering the go or grow phenomenon [Hatzikirou et al., 2010] (one of the key points that regulates the evolution of GB). Obtaining an ABM from this continuum model is important for further exploration of the effect of microenvironment features onto GB progression, such as the role of the immune system, the effect of the composition and structure of the extracellular matrix (ECM) or the growth or formation of new vascular networks (angiogenesis and vasculogenesis, respectively).

In summary, the aim of this work is to obtain mathematical relationships between a broad class of continuum models, that describes the combined effects of reaction, convection, and diffusion, and a general class of ABMs, with an archetypal modelling structure for cell proliferation, differentiation and migration. Then, we apply this methodology to obtain the parameters that regulate the evolution of GB cells in a microfluidic device, able to reproduce important structures such as pseudopalisade and necrotic core formation[Ayuso et al., 2016, 2017].

The work is structured as follows. First, in the methods section, we describe the mathematical structure of the family of ABMs that we consider and how the parameters of the continuum model are inherited by the ABM. We also tackle the way in which the boundary conditions are considered. Then, we present the example of application, the derivation of an ABM for GB progression in microfluidic devices. In this section, we describe briefly the experimental data that is intended to replicate, and how some components of the model structure can be inherited from a reference continuum model of the scientific literature. Subsequently, we demonstrate to reproduce the different experimental configurations using our derived ABM, namely, the pseudopalisade and the necrotic core formation. We close the paper with a discussion on the *in silico* replication of the experiments, the range of the parameters obtained and further possible extensions of the ABM and with the main conclusions of the work.

## 2 Methods

ABMs are defined by a large number of parameters. This and the high correlation among some of them make the parametric fitting process from a set of experimental data cumbersome and many times inaccurate. To overcome these limitations we try here to obtain some important values of such parameters from equivalent continuum models based on a general class of the reaction-diffusion equation. This allows to avoid the fitting process, when the continuum parameters have been previously calibrated, as well as to easily incorporate the dependencies with the different field variables.

In this section, we define the fundamental equations of a general class of ABMs, the methodology proposed for parameter identification, and how the interaction of the agents with the domain walls can be modeled, in an analogous manner to the boundary conditions in PDE-based continuum problems.

### 2.1 Agent-Based Model

The family of ABM considered is the one implemented in PhysiCell [Ghaffarizadeh et al., 2018]. This software is a library for the simulation of ABMs, with special emphasis on cellular applications. It is open source, programmed in C++ and does not require any meshing. Cells are simulated as sphere, being therefore a centered-based ABM approach. To model the species of the microenvironment in which cells move, PhysiCell uses the finite volume method with a software called BioFVM [Ghaffarizadeh et al., 2015].

This class of models considers *n* different cell populations, together with *m* chemical species (signals, nutrients, etc.). Cells can proliferate, migrate, die, and consume and secrete substances. Next, we discuss each of these phenomena in detail, adopting some additional hypotheses to adapt the general formulation in Physicell to some common assumptions from existing scientific evidence.

#### Cell proliferation

Proliferation is modeled by a set of transition probabilities between the *M* different phases of the cell cycle {*X*_1_, *· · · X*_*M*_ }. Each cell *k* is identified by a specific phenotypic phase 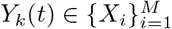. In the absence of the possibility of experimentally measuring all these transitions, living cells are considered to have only one phase *X*_0_. Therefore, we will only have one transition ratio associated to the *i*-th cell population *g*_*i*_, with *i* = 1, *· · ·, n*.

Cell proliferation is assumed to be a Poisson point process so, a cell’s probability of leaving the phase (*X*_0_) and duplicate from a time *t* to a time *t* + Δ*t*, for each cell population, is:

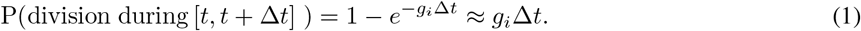

That is, we consider that cell division is memoryless. When cells complete their cycle and divide, there is a duplication of the cells, including the parameters and their associated states.

#### Cell death

Cell death is modeled in a similar way to proliferation, using phases of death and associating probabilities with the change between phases. Cell death occurs by two processes: apoptosis and necrosis. We have modeled both processes with one single transition ratio (*d*_*i*_), between a living cells phase (*X*_0_) and a death phase (*X*_*S*_):

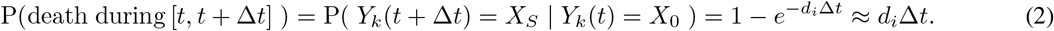

When cells die, a swelling process takes place, in which the cell volume grows and reaches a specific maximal volume Ghaffarizadeh et al. [2018].

#### Cell migration

The velocity of cell movement is defined from a balance of forces which has three contributions: mechanically-driven, random (pedesis) and chemotactically-driven migration. In our example, mechanical forces are not taken into account. Cell velocity is then determined by:

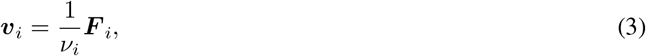

where ***v***_*i*_ is the velocity of each cell of the *i*-th population and *ν*_*i*_ is the drag coefficient for that population. Cell migration, both random (pedesis) and biased (chemotactic) is represented through a force:

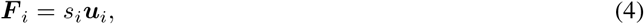

where *s*_*i*_ is the force intensity and ***u***_*i*_ is a unitary vector that is calculated as:

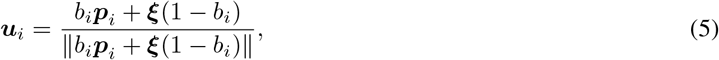

where ***ξ*** is a unit vector with random direction and ***p***_*i*_ is a unit vector indicating the direction of biased migration (for example, oriented towards the gradient of a substance in the microenvironment). Finally *b*_*i*_ *∈* [0, 1] is a bias parameter. If *b*_*i*_ = 0 the motion is totally random, and if *b*_*i*_ = 1, it is totally deterministic and biased along ***p***_*i*_.

#### Species evolution

The chemical species or substrates are considered to be regulating substances in the cell microenvironment, since they are important factors in cell dynamics (particularly when using ABMs for modelling the development of some pathologies such as cancer). The transport equation that defines how the species concentration *S*_*j*_, *j* = 1, *· · ·, m*, varies, is as follows:

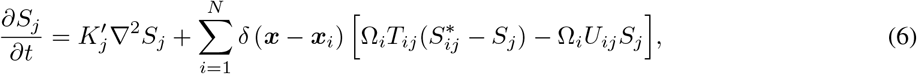

where 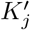 is the species diffusion coefficient, ***x*** is the position vector, ***x***_*i*_ is the position vector of the *i*-th cell position center, Ω_*i*_ is the volume of cell *i, T*_*ij*_ the secretion rate of each cell *i* for each species *j*, 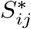 the secretion saturation for each cell of the population *i* of the species *j, U*_*ij*_ is the uptake rate of each cell *i* for the species *j, N* is the total number of cells, and *δ* is the Dirac delta.

### 2.2 Deriving ABM parameters from a continuum model

#### 2.2.1 Model parameters

In this section we show how the parameters of the ABM and those of the continuum model relate to each other.

In particular, we need to establish the explicit expression of the following model functions which in general may depend on the cell population concentrations (*C*_1_, *C*_2_, …, *C*_*n*_), the species concentrations (*S*_1_, *S*_2_, …, *S*_*m*_) and their derivatives. Hence:

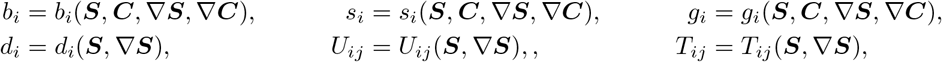

with *i* = 1, …, *n, j* = 1, …, *m*, and:

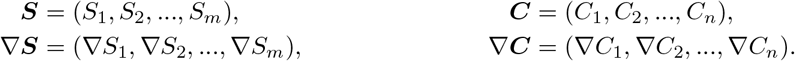

We are interested in deriving these functions from a generic continuum model, based on transport equations, that incorporates the same phenomena (migration, proliferation, death, consumption and secretion), together with the chemical species diffusion as moderating agents in the microenvironment. The fundamental equation of the continuum model for each population *C*_*i*_ is the Reaction Convection Diffusion Equation (RCDE), written as:

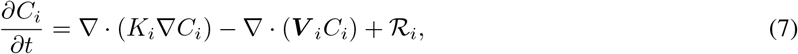

where *K*_*i*_ is the diffusion coefficient, ***V*** _*i*_ the drift velocity and ℛ_*i*_ the reaction term, that can be decomposed in a source and a sink term:

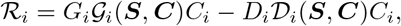

where the *G*_*i*_ and *D*_*i*_ are the growth and death characteristic rates and 𝒢_*i*_ and 𝒟_*i*_ are context dependent functions accounting for cell behavior, that may depend on species or nutrient availability and cell concentration.

Besides, the fundamental equation for each chemical species in the microenvironment *S*_*j*_ writes:

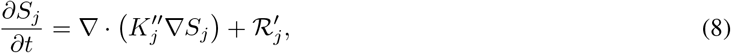

where for each species, 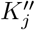 is the diffusion coefficient and 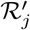 is the reaction term, which can be written in terms of a production and an consumption or intake term as:

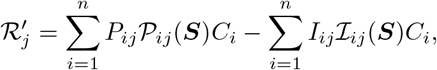

where *P*_*ij*_ and *I*_*ij*_ are the production and consumption characteristic rates of the species *j* by the cell *i* and 𝒫_*ij*_ and ℐ_*ij*_ are again context dependent functions.

In what follows we present the relationships obtained between the parameters and functions in both models. For the mathematical details of how the following relationships have been obtained, the reader is referred to Appendix A.

##### Cell migration

Using a random walk model it is possible to derive mathematical equations relating the values of *s*_*i*_ and *b*_*i*_ to the advection-diffusion components of the continuum model. According to Eqs. (44), (45) and (46), the final relationships obtained for the *i*-th cell are:

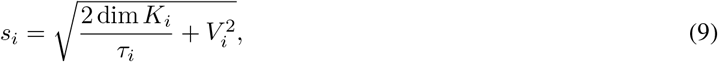

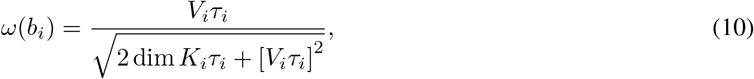

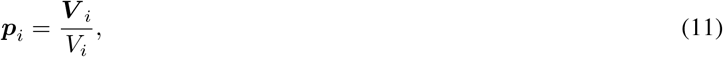

where, for each population, *V*_*i*_ is the modulus of the convective velocity, dim is the dimension of the space, *τ*_*i*_ is the persistence time, and *ω* is a function that defines how random the movement is, given by

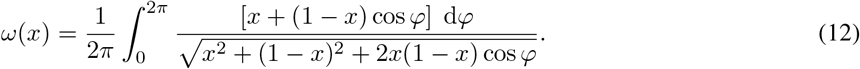

##### Cell proliferation and death

As explained above, the formulation of cell proliferation of the ABM includes one parameter for each cell population (*g*_*i*_). If we compare the proliferation terms of both models, according to Eq. (48) we obtain the relation:

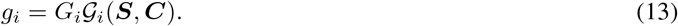

Besides, cell death in the ABM is regulated by one parameter (*d*_*i*_). The relationship between the ABM death ratio and the sink term in the continuum model is, according to Eq. (49):

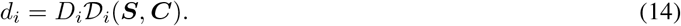

##### Species evolution

We consider that the diffusion process is homogeneous, and thus 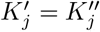. Species uptake and secretion in the ABM is regulated by the parameters *U*_*ij*_, *T*_*ij*_ and 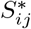, as we saw in Eq. (6). For the case where Ω_*i*_, 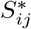 and *U*_*ij*_ are equal for all the cells of the same population, by comparing both models, we get the following from Eq.(53)

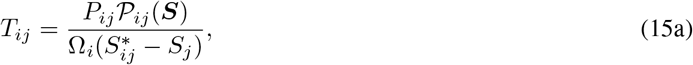

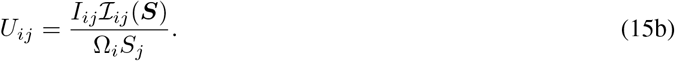

#### 2.2.2 Operating conditions

Usually, when using ABM for recreating experimental data, initial and boundary conditions play an important role. We calculate the initial number of agents of population *i* (*N*_*i*,0_) as:

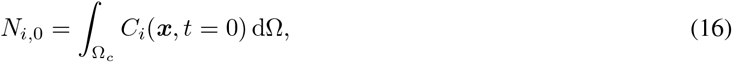

where Ω_c_ is the volume of the chamber of the microfluidic device, and *C*_*i*_(***x***, *t* = 0) the initial cell concentration used in the experiment. Then, we distribute the obtained number of cells randomly in the domain, so that they follow the same distribution than the experimental data.

With respect to the boundary conditions, it shall be noted that domains much larger than the region of interest are typically used in ABMs to prevent the contours from affecting the behavior of the agents. However, for certain experimental configurations, it is not possible to neglect the impact of the boundaries of the domain, since the agents are in a confined environment and their behavior close to the boundaries is key because at these locations is at which the supply of species, e.g. nutrients, occurs. Besides, while a potential function can be activated at the agent level for preventing them from escaping the domain, we decided to model this interaction explicitly to control the agent behavior close to the boundaries. That said, the limits of the computational domain can be divided into two different regions depending on their behaviour: impenetrable barriers for the agents or permeable membranes that allow agents pass through them, with greater or lesser difficulty.

Hence, the following condition is defined to prevent agents from escaping the domain through the chamber walls, mimicking no flux Neumann boundary conditions in continuum models: if the agents are in contact with the wall, that is, they are at a distance from the wall lower than their radius, and at the instant *t* the velocity component perpendicular to the wall is directed outward from the domain (***v***_*t*_*·****n*** *<* 0), then their velocity is modified at the instant *t* + 1 so that the direction of the component perpendicular to the wall is inverted emulating a collision:

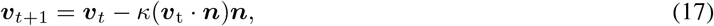

where ***n*** is the normal vector to the wall, and *κ >* 0 depends on whether the collision is elastic or not. From Eq. (17) we observe that

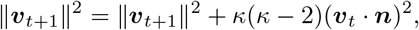

Therefore, for an elastic collision, *κ* = 2. If *κ >* 2, the boundary is transferring energy to the agent and if 0*≤ κ* 2*≤* the collision is inelastic, being perfectly inelastic when *κ* = 1.

However, for permeable membranes, the agents may escape from the domain by crossing the membrane. To include this leakage phenomenon, we define a probability *q* = P(crosses during [*t*, Δ*t*]). This probability is added to the previous condition to determine whether or not the agent velocity is modified when the agents approach the walls of the domain in contact with the channels. For each agent approaching the domain edge, the agent escapes with probability *q*. In particular, if *q* = 0 the condition defined in Eq. (17) is applied to every agent, while for *q* = 1 we allow that every agent arriving to the wall escapes. We assume that the probability *q* of agent escaping is homogeneous throughout each boundary region and along time.

In summary, for the operating conditions we have two parameters (*κ* and *q*) that depend on the characteristics of the experiment.

## 3 Application to glioblastoma progression in microfluidic devices

### 3.1 *In-vitro* experiments

Cell culture in microfluidic devices has demonstrated to be able to reproduce the main features of GB in physiological conditions, as are the formation of necrotic core [Ayuso et al., 2016] and of migrating structures called pseudopalisades [Ayuso et al., 2017].

The experiments were performed in microfluidic devices with a gas-impermeable central chamber, made of Cyclic Olefin Polymer (COP), and two side microchannels. Cells of the commercial GB line U251-MG embedded in a collagen hydrogel were seeded homogeneously into the central chamber, and culture medium with oxygen was perfused through the side channels [Ayuso et al., 2016, 2017]. A first experiment was performed in a microfluidic system with a 916 *µ*m wide chamber, and low initial cell concentration (4*·*10^6^ cell*/*mL) [Ayuso et al., 2017] reproducing the formation of a pseudopalisade; while a second experiment was performed in a device with a 2000 *µ*m wide chamber, and high initial cell concentration (40*·*10^6^ cell*/*mL) [Ayuso et al., 2016], this time reproducing the formation of a necrotic core. To trigger the pseudopalisade formation, one of the channels was closed, preventing oxygen from flowing through it and generating a gradient in the chamber, while for the necrotic core formation oxygen-rich medium was perfused through both channels. The area of the chamber that is not in contact with the channels acts as a barrier for cells, that cannot escape from the chamber. On the other hand, the lateral channels, allow oxygen supply, but provide the cells with an escape route (see figure 1 for a schematic representation of the experimental configuration). From these experiments we obtained fluorescence images of alive and dead cell concentration at the initial (0), third and sixth days.

**Figure 1:**
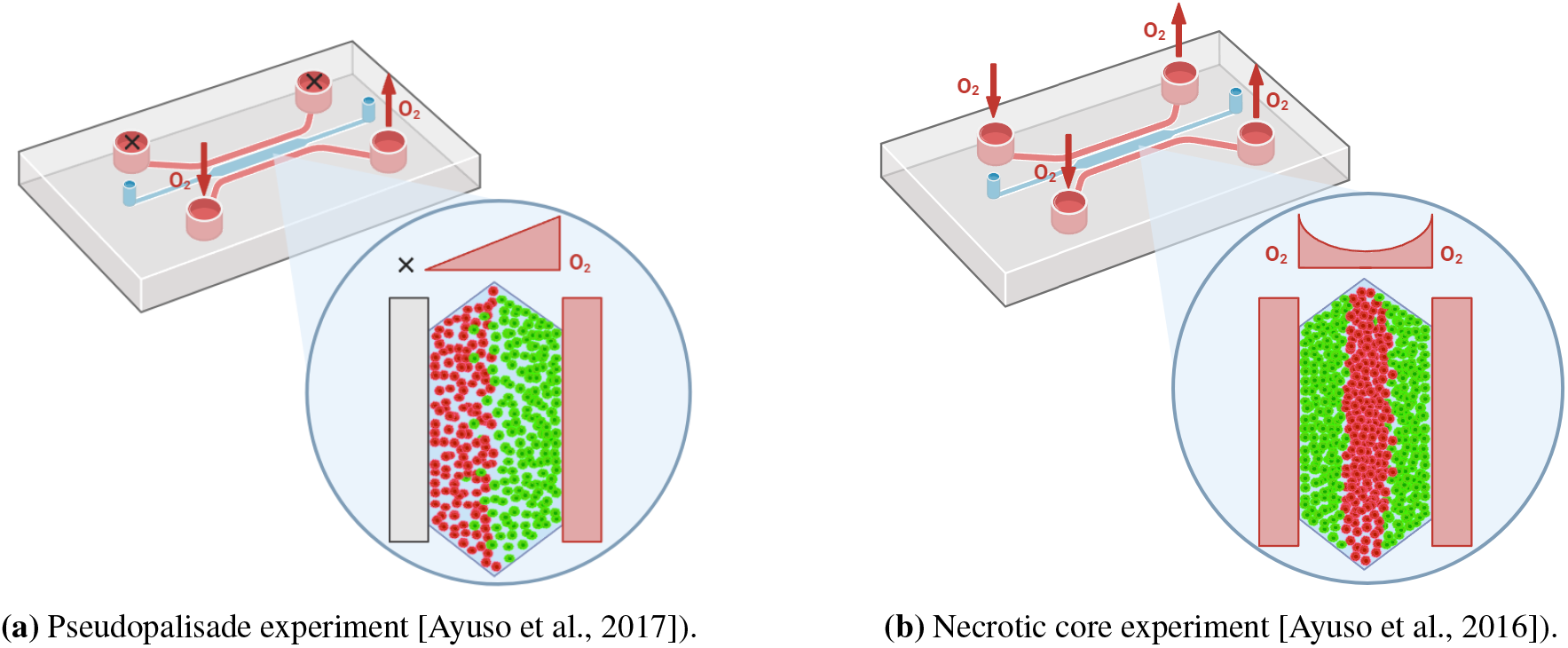
Schematic representation of the experimental configuration of the pseudopalisade and the necrotic core. Green cells represent alive GB cells, and red cells represent the dead GB cells. In the necrotic core experiment, oxygen medium is perfused through both lateral channels. In the pseudopalisade experiment, only one of the channels is perfused with oxygen. Created with BioRender.com.

### 3.2 Particularities of glioblastoma evolution

Research on the origin of pseudopalisades formation and necrotic zones has shown that hypoxia plays a key role in this process [Brat and Van Meir, 2004]. In this section, we particularise the general ABM from the previous section to include the impact of hypoxia on GB evolution. Hence, we made the following assumptions:

We consider only one cell population of tumor live cells, all of which have the same phenotype (not considering the heterogeneity of the GB tumor) and that cells do not secrete any chemical species that affect the behaviour of the GB, so we do not take into account the cell secretion. We also consider dead cells, and oxygen as the moderator environment substance.

As was demonstrated in previous experiments and mathematical models [Ayuso et al., 2016, 2017, Ayensa-Jiménez et al., 2020], cell behavior (migration, proliferation, death and uptake) is regulated by oxygen since, as commented in the introduction, it is the chemical species that mainly regulates the formation of the pseudopalisades and necrotic core. In particular:

- Migration and proliferation are oxygen-dependent according to the go or grow model [Hatzikirou et al., 2010]. That is, cells migrate more for low oxygen levels and, the higher the oxygen level is, the more the cells proliferate. The metabolic switch between migratory and proliferative metabolic activity occurs at a certain oxygen threshold.
- The death process depends on the oxygen concentration via anoxia-induced necrosis [Steinbach et al., 2003]. The dead cells are considered as inert so they cannot proliferate, migrate or consume oxygen.
- Oxygen consumption depends on the oxygen level, as it is common in many biochemical processes of which oxidative phosphorylation is an example: the more oxygen there is in the environment, the greater the cellular consumption of oxygen is [Martínez-González et al., 2012].

Besides, migration and proliferation also depend on the cell concentration. Cells can only migrate or proliferate when there is sufficient space around them (that is, when the surrounding tissue is not cell saturated) [Stramer and Mayor, 2016].

From these assumptions and simplifications, we derive the functional dependencies of the GB on chip ABM parameters with the parameters of a GB on chip continuum model. As mentioned in the introduction (Section 1), we chose the model described in Ayensa-Jiménez et al. [2020] model as reference. This model simulates GB evolution under hypoxic conditions in microfluidic devices using reaction-diffusion partial differential equations for describing the evolution of the concentration of alive (*C*_1_ = *C*_n_) and dead (*C*_2_ = *C*_d_) cells as well as of oxygen (*S*_1_ = *S*). It incorporates two parameters related to migration phenomena (the chemotaxis coefficient, *χ*_n_, and the diffusion coefficient, *K*_n_), one parameter related to proliferation (the characteristic proliferation time, *τ*_g_), another to death (the characteristic death time, *τ*_d_), one parameter related to uptake (consumption rate, *α*), one parameter that controls the cell saturation (cell saturation concentration, *C*_sat_), four metabolic parameters related to the effect of oxygen on the different phenomena (the Michaelis-Menten constant, *S*_M_, the hypoxic threshold *S*_H_, the anoxia threshold *S*_A_ and the sensitivity to anoxia Δ*S*_A_) and the oxygen diffusion coefficient *K*^*′*^. The results of the derivations for the different phenomena are detailed next.

#### Migration

This phenomena is defined in the ABM through *b*_1_ = *b*_1_(*S, ∇ S, C*_n_), *s*_1_ = *s*_1_(*S, ∇ S, C*_n_). Note that *s*_2_ = 0 since dead cells do not have migratory capacity. The functions relating *b*_1_, *s*_1_ with the oxygen and cell concentration are monotone decreasing with respect to both *S* and *C*_n_.

If we particularize Eqs. (9) and (10) to the migration model in Ayensa-Jiménez et al. [2020], we get:

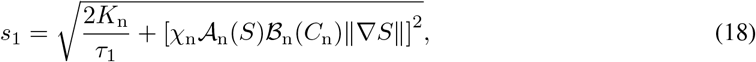

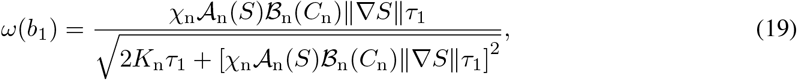

where *τ*_1_ is the migration persistence time of live cells, that is the time that cells maintain their migration direction (identified with the random walk marching time), *ω* is the function given by Eq. (12), and 𝒜_n_ is an activation function that regulates migration as a function of oxygen level (*S*) and the hypoxic threshold (*S*_H_), incorporating the go or grow [Ayensa-Jiménez et al., 2020]:

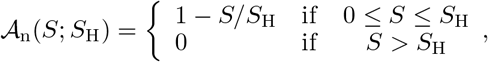

and ℬ_n_ is an activation function that regulates migration as a function of cell concentration (*C*_n_) and cell saturation concentration (*C*_sat_) [Ayensa-Jiménez et al., 2020]:

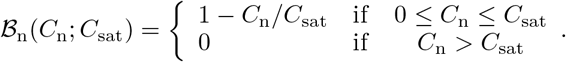

#### Proliferation

Cell growth is regulated by the expression of the growth rate *g*_1_ = *g*_1_(*S, C*_n_). Note that *g*_2_ = 0 since dead cells are inert and cannot proliferate. The function relating *g*_1_ with the oxygen and cell concentration is monotone decreasing with respect to *C*_n_ and monotone increasing with respect to *S*. Considering only one cell phenotype and the model in Ayensa-Jiménez et al. [2020], Eq. (13) becomes:

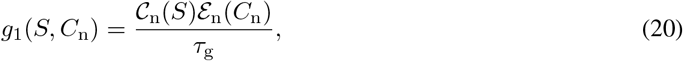

where 𝒞_n_ is an activation function, which states, according again to the go or grow hypothesis, that cells proliferate differently depending on the oxygen level, being this behavior regulated by the hypoxic threshold (*S*_H_):

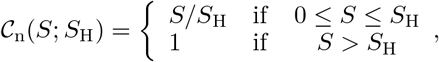

Besides, ℰ_n_ is a saturation function, which implies a logistic growth setting a limit on the cell concentration:

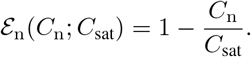

#### Death

The function regulating death in the ABM is *d*_1_ = *d*_1_(*S*). The function is monotone decreasing with respect to *S*. Of course, *g*_2_ = 0. The continuum model has one parameter associated to death (the characteristic death time, *τ*_d_), so Eq. (14) becomes:

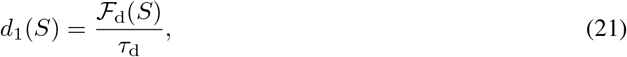

where *ℱ*_d_(*S*) is a hyperbolic tangent which defines the dependence of death on the oxygen level (*S*):

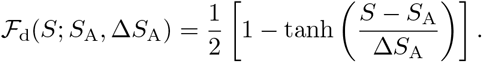

#### Uptake

For this specific case, as there is no oxygen production and only live cells consume oxygen, we have *P*_11_ = *P*_21_ = 0 and *I*_11_ ≠ 0, *I*_21_ = 0. In addition, ℐ_11_(*S*) is defined in the continuum model in terms of two parameters related to the uptake (the consumption rate, *α*, and the Michaelis-Menten constant, *S*_M_).

Comparing the consumption terms between the continuum and ABM, Eq. (15b) rewrites:

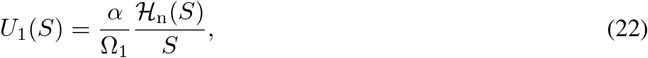

where *ℋ*_n_(*S*) is the Hill function:

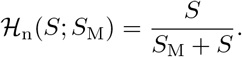

#### Operating conditions

As discussed in section 3.1 it is necessary to define the initial and boundary conditions for the experiments of GB progression in microfluidic devices.

Usually, in the microfluidic devices, the chamber edges are demarcated into two distinct regions: one that does not interface with the channels, and another that is in direct contact with the channels.

The walls with no contact with the supply channels are considered as impermeable to oxygen and cells so following the discussion of section 3.1, we consider for this boundaries *q* = 0 and *κ* = 2. However, it is an experimental observation that some cells reach the side channels, so the interaction cell-channel membrane is more complex. For each particular experiment, the parameters *κ* and *q* are considered as fixed although unknown and had to be fitted.

Regarding the chemical species, we also consider the two options. For the chamber-channel interface, the oxygen level is considered constant and known, therefore imposing a Dirichlet condition in the domain). In the case of the sealed walls of the microfluidic device, a no flux condition is imposed, that is, a Neumann boundary conditions).

### 3.3 Results

Based on the functions and parameter values obtained in Section 3.2 as well as the initial and boundary conditions described in Section 2.2.2, we performed the simulations of the GB experiments described in section 3.1. The size of the chamber used in the 2D pseudopalisade simulation is the same as the one used for the GB experiment (with width *L*_*x*_ = 916 *µ*m and lenght *L*_*y*_ = 916 *µ*m). For the necrotic core simulation, we used a smaller section (*L*_*x*_ = 2000 *µ*m and *L*_*y*_ = 200 *µ*m) of the total chamber (*L*_*x*_ = 2000 *µ*m and *L*_*y*_ = 3000 *µ*m), since the behavior is homogeneous along the y-axis, and to reduce the computational cost.

In order to establish the cellular volume, we searched literature on GB cell size, and have found different sizes for GB primary cells and GB cell lines. For example, for the cell line U87, in the database of BioNumbers [Milo et al., 2009] the cell size varies from 12-14 *µ*m. Grimes et al. [2016] uses a method to estimate the mass/volume of different cell lines, obtaining a 12.5 *µ*m for U87 cell diameter. However, in [Oraiopoulou et al., 2017] they estimated from in vitro experiments the mean cell diameter for GB cell lines from 3 different patients, obtaining 15, 16 and 19 *µ*m respectively, and 21.5 *µ*m for the U87 cell line. In [Jenner et al., 2022] they assume a cell diameter of 21.5 *µ*m for simulate the GB cells. As this parameter has no impact on cell behavior, influencing only the value of cell uptake (as it depends on cell volume, see Eq.22.), we decide to keep the default cell diameter imposed in PhysiCell (16.8 *µ*m) [Ghaffarizadeh et al., 2018], as it is similar to an average radius of all values found in the literature.

According to literature, the persistence time (*τ*_1_) on cancer migration go from 3 to 30 min [Weiger et al., 2010], depending on the experimental conditions. In [Ghaffarizadeh et al., 2018] for breast cancer they use 15 min. And in [Jenner et al., 2022] the results show that for most of the GB cells the persistence time is between 20 and 30 minutes. So we decided to impose a persistence time of 20 minutes to reproduce the *in vitro* results.

As stated in the previous section, the behaviour of cells is determined by the following non-linear functions:

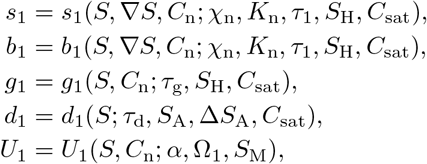

which are defined (explicitly or implicitly) in Eqs. (18), (19), (20), (21) and (22)) respectively, depend on several parameters whose value is defined in Table 1, together with the oxygen diffusion coefficient.

**Table 1:** ABM parameters and functions for simulating glioblastoma progression. The specific value of the parameter is displayed when possible, and for parameters that depend on other variables, the equation in text where the formula in detailed is referenced.

| Parameters | Symbol | Value/Expression | Units |
| --- | --- | --- | --- |
| Oxygen diffusion coefficient | $K'$ | $6 \times 10^4$ | $\mu\text{m}^2/\text{min}$ |
| Cell diffusion coefficient | $K_n$ | 3.0 | $\mu\text{m}^2/\text{min}$ |
| Cell chemotaxis coefficient | $\chi_n$ | 45 | $\mu\text{m}^2/(\text{min} \cdot \text{mmHg})$ |
| Cell volume | $\Omega$ | 2494 | $\mu\text{m}^3$ |
| Persistence time | $\tau_1$ | 20 | min |
| Proliferation characteristic time | $\tau_g$ | $12 \times 10^3$ | min |
| Hypoxic threshold | $S_H$ | 7 | mmHg |
| Cell saturation concentration | $C_{\text{sat}}$ | $5 \times 10^{-5}$ | cell/$\mu\text{m}^3$ |
| Death characteristic time | $\tau_d$ | $2.88 \times 10^3$ | min |
| Anoxia threshold | $S_A$ | 1.6 | mmHg |
| Sensitivity to anoxia | $\Delta S_A$ | 0.1 | mmHg |
| Cell uptake coefficient | $\alpha$ | $6 \times 10^4$ | $(\text{mmHg} \cdot \mu\text{m}^3)/(\text{cell} \cdot \text{min})$ |
| Michaelis-Menten constant | $S_M$ | 2.5 | mmHg |

The parameters related to the changes in cellular volume (such as the fluid fraction of a cell, the cell nuclear volume, the solid cytoplasmic volume, among others), as they have no impact for our results, beyond visualization, have been kept at their reference value as defined in Ghaffarizadeh et al. [2018].

Regarding oxygen, in the necrotic core we consider that the oxygen concentration supplied through both channels is constant and known (imposing a Dirichlet condition in the domain, whose values are indicated in Figure 2), and for the pseudopalisade we consider that there is a sealed channel, and therefore a no flow condition is imposed (Neumann boundary condition). A scheme of both configurations can be seen in Figure 2. The initial oxygen concentration is homogeneous throughout the chamber (7 mmHg for the necrotic core experiment and 2 mmHg for the pseudopalisade experiment [Ayensa-Jiménez et al., 2020]).

**Figure 2:**
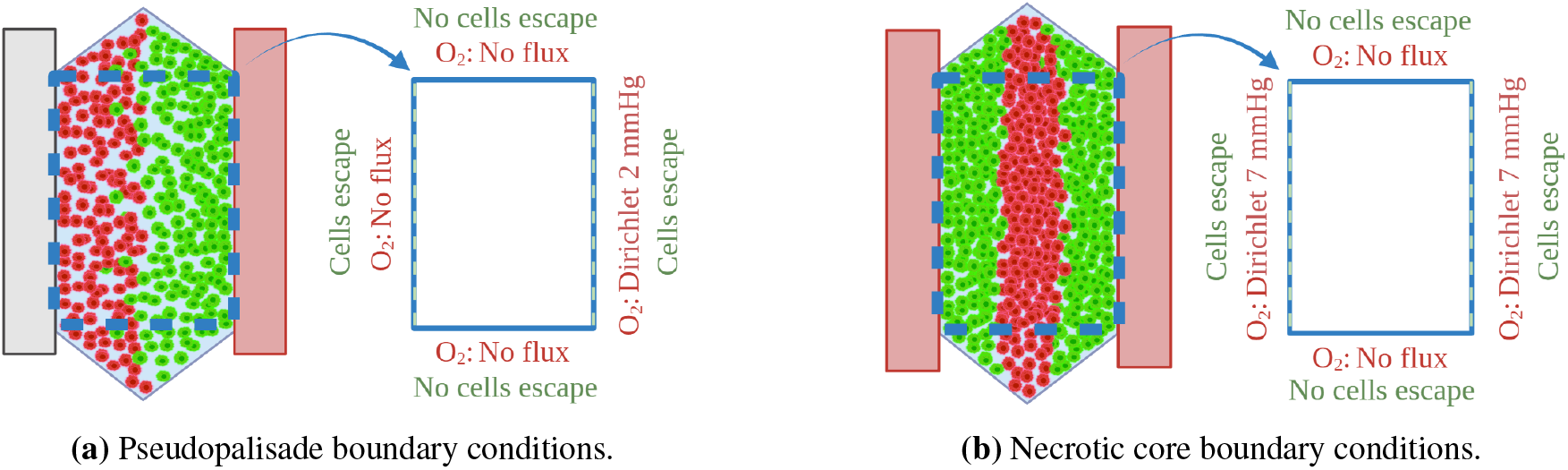
Schematic representation of the chamber, and the operating conditions used for the simulation of the pseudopalisade and necrotic core. The boundaries of the chamber that are in contact with the channels (dashed line) allow cells to escape. Created with BioRender.com.

For the operating conditions, the probability used to reproduce the experimental results is: *q* = 1 for the case of necrotic core and *q* = 0.15 for the case of the pseudopalisade. These values have been adjusted using the objective cost function

(*T*) that minimises the differences between ABM and experimental results computed over an horizontal central line perpendicular to lateral channels:

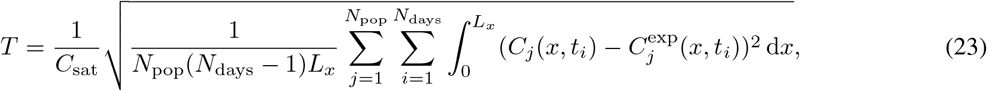

where *N*_pop_ is the number of populations (alive, dead) for the experiment, *N*_days_ the number of days for the experiment, *C*_*j*_ is the ABM cell concentration and 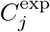 is the experimental cell concentration for the experiment.

Given the stochastic nature inherent in the phenomena driving cell evolution within the model, cells exhibit varied responses when subjected to identical local environmental conditions. This variability underscores the necessity of conducting numerous replications of simulations to ascertain population behaviors and draw conclusive insights. In our study, we conducted 30 replicates for each simulation, ensuring convergence of the mean cell concentration to a constant value.

#### 3.3.1 Pseudopalisade

Figure 3 displays the position of the cells for the initial, third and sixth days in one representative replicate. Green cells represent living cells while red cells are dead cells. The cells are initially distributed to replicate the experimental initial concentration. When the simulation starts, cells begin to die in the areas with less oxygen (left region of the chamber), and those that survive migrate to the right area, which is more oxygenated.

**Figure 3:**
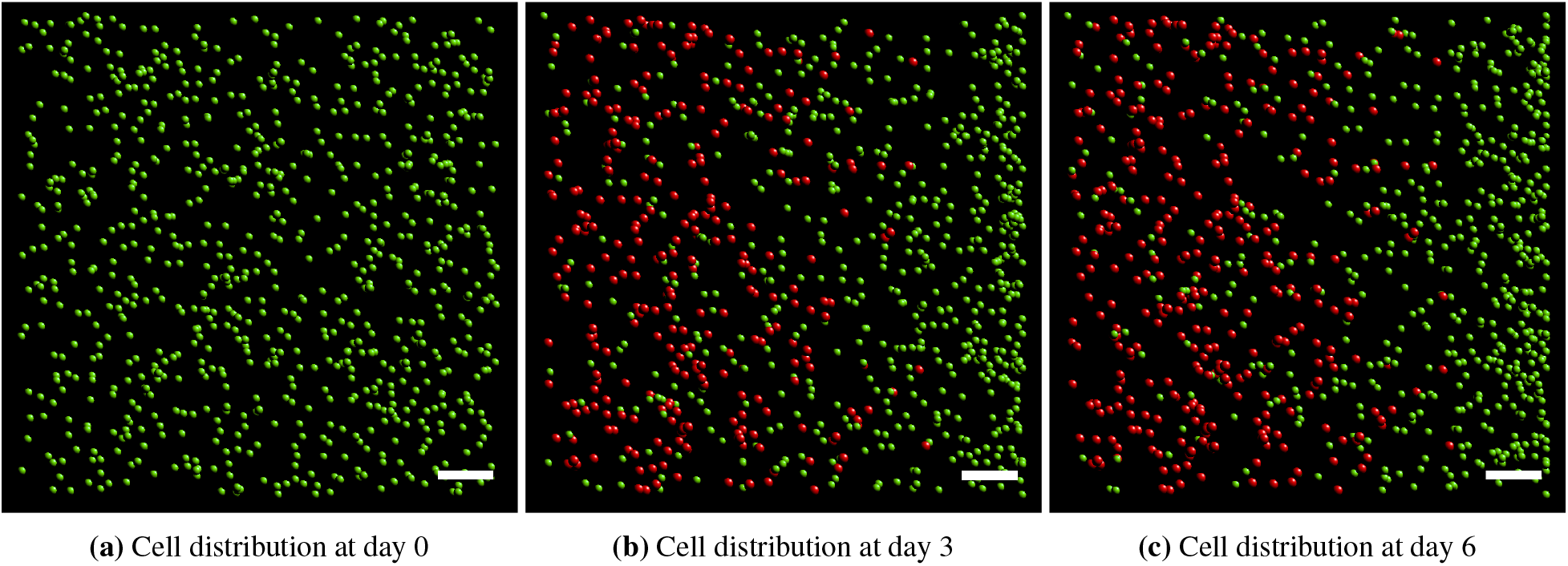
Pseudopalisade results. Initial, medium and final states (alive or dead) of the cells in the ABM. Alive cells are represented with green spheres while dead cells are represented with red spheres. Scale bar: 100 *µ*m

If we now plot the mean cell concentration of the replicates with the 95% confidence interval and compare it to both the experimental results and to the results obtained with the continuum model, we obtain Figure 4. The figures presented demonstrate concordance between the experimental and simulated outcomes, with a value of the objective cost function for the ABM of *T* = 0.0084. The value of *T* for the continuum model and the experimental data is *T* = 0.0106 in Ayensa-Jiménez et al. [2020], so we achieved an improvement in the results.

**Figure 4:**
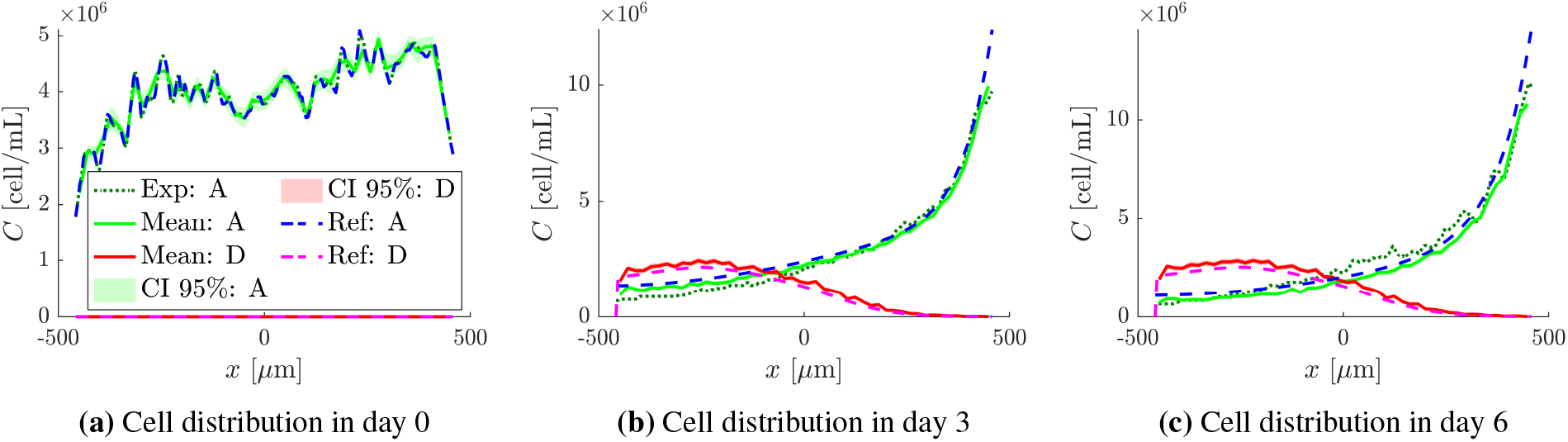
Comparison between experimental and simulated results of the pseudopalisade formation. The experimental results for alive cells (there were no data on the dead cells for this experiment) are displayed with a dark green dashed line, together with the median and 95% Confidence Interval (CI) of the results produced by the ABM for both alive (green) and dead (red) cells, and the continuum results of alive (blue dashed line) and dead (magenta dashed line) cells. Ref: Ayensa-Jiménez et al. [2020] with the parameters of Table 1.

This results will be discussed in more detail in the next section.

#### 3.3.2 Necrotic core

Analogously to the pseudopalisade simulation, the results of cell distribution for the simulation of the necrotic core formation are displayed in Figure 5. This figure shows a simulation of the complete chamber, although as previously mentioned, the graphs in Figure 6 are obtained from a smaller section of the chamber in order to reduce the computational cost.

**Figure 5:**
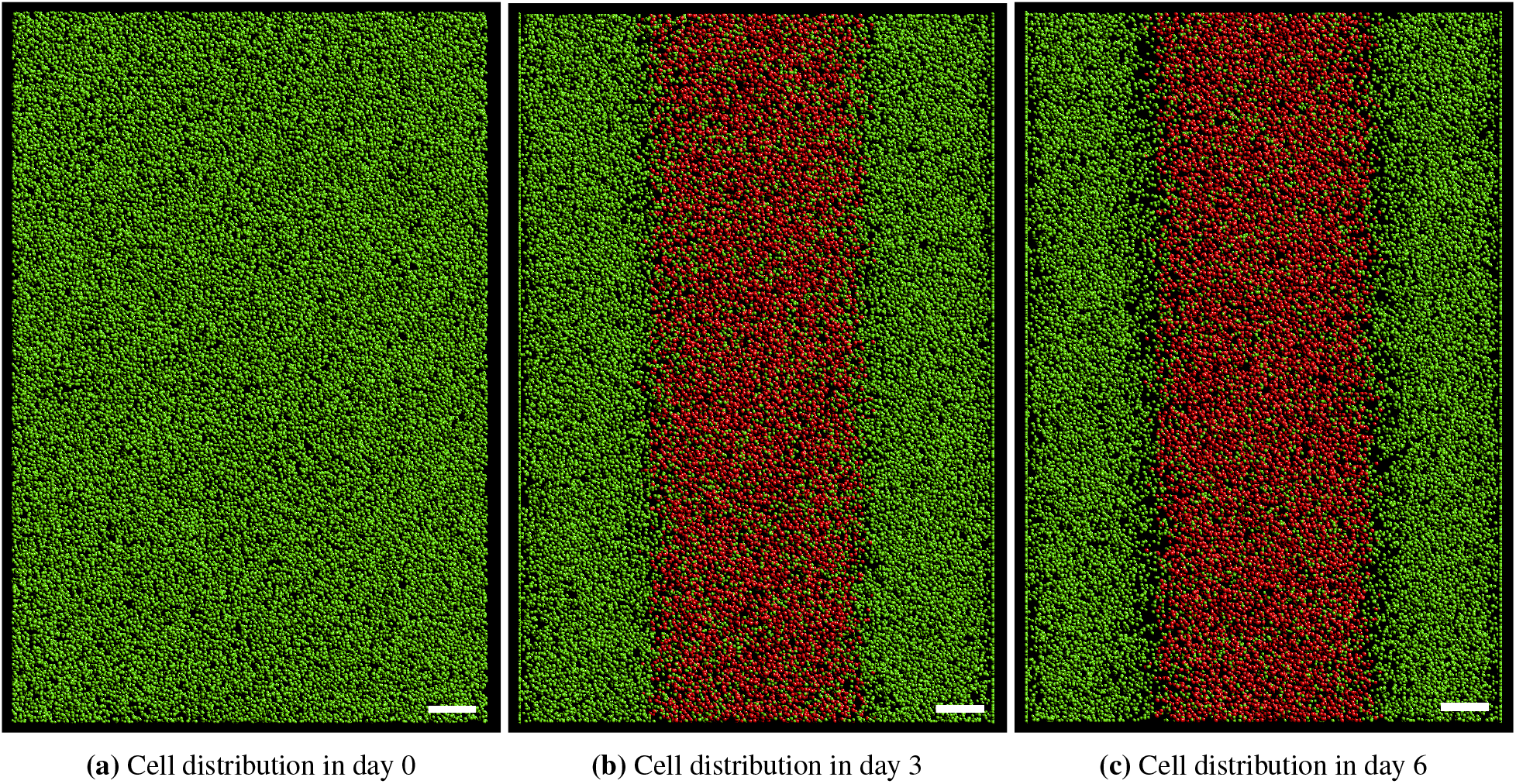
Necrotic core results. Initial, medium and final state of the cells in the ABM. Scale bar: 200 *µm*

**Figure 6:**
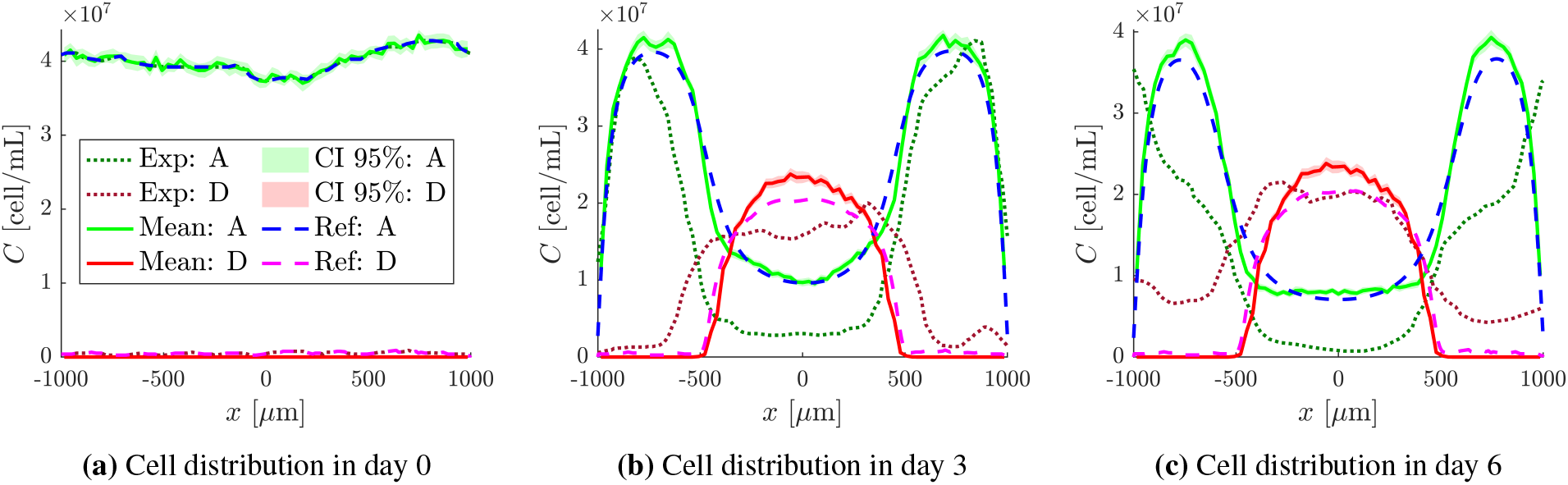
Comparison between experimental and simulated results of the necrotic core formation. The experimental results for alive cells are displayed with a dark green dashed line, and the dead cells with a garnet dashed line, together with the median and 95% Confidence Interval (CI) of the results produced by the ABM for both alive (green) and dead (red) cells, and the continuum results of alive (blue dashed line) and dead (magenta dashed line) cells. Ref: Ayensa-Jiménez et al. [2020] with the parameters of Table 1.

The simulation starts with the same distribution of alive cells as the corresponding experiment. Over time, given the high cell density, which induces an oxygen-deprived area in the central region of the chamber, and the continuous supply of oxygen through the lateral channels, cell migration is notably directed towards the lateral channels, which are more oxygenated (see Figure 5c). Within the central region of the chamber, a decline in oxygen concentration due to cellular uptake leads to cell death.

In Figure 6 the comparison between experimental, continuum and ABM results is represented. The computed results exhibit both qualitative parity and quantitative resemblance to the continuum results. Comparing the results of the ABM to the experimental results, we obtain *T* = 0.1706, which is close to the value obtained between the continuum model and the experimental data, *T* = 0.1723 in Ayensa-Jiménez et al. [2020]. However, notable disparities are evident in the experimental live cell profile, particularly at the central region of the chamber, and in the dead cell profile at the boundaries.

## 4 Discussion

Modeling a problem as complex as the evolution of cancer is not easy. The many coupled phenomena acting in this biological process make it impossible to obtain experimental results fully controlling all conditions. In addition, ABMs, being highly parametric, require more information, at the cellular and environmental levels, than other types of models, such as continuum ones, for parameter calibration. That is why, in this work, we used a continuum model already calibrated [Ayensa-Jiménez et al., 2020], using the information obtained in that work to derive the parameters of the ABM analytically, avoiding to directly perform the fitting on the ABM, that is reduced here to one single operation parameter, *q*. Instead, mathematical relations have been used to define the value of the parameters and functions in the ABM related to the continuum model parameters.

For this purpose, each phenomenon involved in the evolution of this tumor has been isolated. With the mathematical deductions obtained, it is possible to bridge the gap between both models, allowing to work with them simultaneously and take advantage of the benefits of each of them.

The resulting ABM can reproduce the evolution of GB obtaining the same results obtained with the continuum model, replicating the main histopathological GB features (the formation of necrotic cores and pseudopalisades) recreated in microfluidic devices, but with some additional advantages. First, the ABM not only reproduces the continuum model but also incorporates inherent random effects typical of biological models, providing a more natural explanation of the biological processes. This is due to the stochastic nature of some of the processes as well as of the initial cell distribution which result in different results for the different replicates carried out. Moreover, additional relevant phenomena can be easily incorporated into this model, such as the mechanical interaction between cells or between cells and the environment. The methodology presented herein, allowed us to obtain a first GB ABM, which can be further refined at the individual level thanks to the benefits of ABMs and incorporate aspects that do not have a straightforward implementation in continuum models.

In relation to the parameters, an analysis of the range of possible values has been carried out to see if they are within the ranges identified in the literature. Many of the parameters (Table 1) used in this model have been directly obtained from Ayensa-Jiménez et al. [2020], so we will therefore focus on the parameters that have a mechanistic interpretation from the biological point of view, that is, *s*_1_, *b*_1_, *g*_1_, *d*_1_ and *U*_1_, that do not appear in the continuum model:

### Migration speed

(*s*_1_). There are two components defining the migration speed: directed (due to chemotaxis) and random (due to pedesis). Considering the case when migration is purely random (for instance, in absence of oxygen gradients), the speed is defined by:

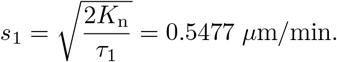

This value is in agreement with what is found in the scientific literature. Indeed, Diao et al. [2019] found that the average migration speed for four GB cell lines range from 0.15 *µ*m*/*min to 0.57 *µ*m*/*min, and they observe the motility of U251 is lower that in other cell lines, with a mean migration speed of (0.17*±*0.02) *µ*m*/*min. Besides, Jenner et al. [2022] use a velocity of 0.1 to 1 *µ*m*/*min on their simulation. Finally, Kaphle et al. [2018] report that the migration speed for different conditions of hydrogel and culture dish vary between 0.3 *µ*m*/*min to 1.04 *µ*m*/*min.

If now we focus on migration dominated by the chemotaxis term, under physiological conditions, the distance between vessels can be estimated as *L∼* 50 *µ*m [Duvernoy et al., 1983, Ndubuizu and LaManna, 2007] and the oxygen level varies from *S∼* 10 mmHg to *S∼* 2 mmHg [Hoffman et al., 1996, Ndubuizu and LaManna, 2007] so ∥ ∇*S* ∥∼ 0.16 mmHg*/µ*m (or using Henry’s law [Popel, 1989], *∥∇S∥ ∼* 2.2 *×* 10^*−*4^ mM*/µ*m). Therefore, assuming that 𝒜_*n*_ and ℬ_*n*_ are bounded decreasing functions, the maximal value of the cell migration velocity is:

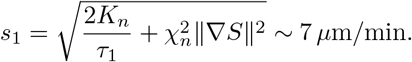

In the work by Agosti et al. [2018a], they found a chemotaxis coefficient in the range 8 – 12 mm^2^ *·* mM^*−*1^*·*d^*−*1^ so the cell velocity is *v*_chemo_ *∼*2 *µ*m*/*min, which is of the order of *s*_1_, taking into account that the latter is an upper bound.

### Bias parameter

(*b*_1_). The movement of the cells depends on the oxygen level and the cell concentration. From Eq. (19) we observe that when *C*_n_*≥ C*_sat_, *S≥ S*_H_ or *∇ S* = **0**, *b*_1_ = 0, that corresponds to a random movement, whereas if ∥∇*S*∥ → ∞, *b*_1_ = 1 that corresponds to a perfect drift. For a fixed value of the oxygen gradient, as 𝒜_*n*_ and ℬ_*n*_ are bounded decreasing functions, the maximal value of *b*_1_ is achieved for low cell concentrations and hypoxic environments, in agreement with the experimental evidence [Brat et al., 2004,

Ayuso et al., 2017, *Monteiro et al., 2017]*. *Under physiological conditions, as* ∥∇*S*∥ ∼0.16 mmHg*/µ*m, *ω* ∼ 0.95, which in turns imply *b*_1_*∼* 0.8, typical of purely chemotaxis-driven systems [Jenner et al., 2022, Ozik et al., 2018]. Hence, the physiological values of the microenvironment translate into reasonable values for *b*_1_.

### Proliferation rate

(*g*_1_). It is straightforward to see from Eq. (20) that 0*≤ g*_1_*≤ g*_max_ with *g*_max_ = 8.3*×* 10^*−*5^ min^*−*1^. This range is in agreement with the values found in the scientific literature, that are very heterogeneous Rockne et al. [2010], Hathout et al. [2016], Agosti et al. [2018a].

### Death rate

(*d*_1_). Similarly, from Eq. (21) it can be seen that 0 *≤ d*_1_*≤ d*_max_ with *d*_max_ = 4.2 *×*10^*−*4^ min^*−*1^. This range is also in agreement with the values found in the scientific literature Frieboes et al. [2007], Martínez-González et al. [2012], Agosti et al. [2018b].

### Cell uptake

(*U*_1_). Regarding cell uptake, it follows from Eq. (22) that Ω_1_*U*_1_(*S*)*S →* 0 when *S →* 0 and Ω_1_*U*_1_(*S*)*S → α* when *S → ∞* so the oxygen consumption by the cell is of the order *α* = 1.0 *×* 10^*−*9^ mmHg *·* mL *·* cell^*−*1^ *·* s^*−*1^, consistent with the values found in the literature Gerlee and Anderson [2007], Daşu et al. [2003]. Moreover, if we consider the unitary uptake, 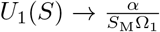 when *S →* 0 and *U*_1_(*S*) *→* 0 when *S → ∞*. Considering a cell of characteristic length *R* = 8.4 *µ*m so that Ω_1_ *∼* 2500 *µ*m^3^ = 2.5 *×* 10^*−*9^ mL and provided that *α* = 2.0 *×* 10^*−*9^ mmHg *·* mL *·* cell^*−*1^ *·* s^*−*1^ and *S*_M_ = 2.5 mmHg we obtain that 0 *≤ U*_1_ *≤* 0.32 cell^*−*1^ *·* s^*−*1^, that is consistent with the values found in the literature Agosti et al. [2018b].

In summary, starting from a continuum model we have been able to define an ABM for the evolution of GB and, at this point, the ABM can be extended to improve the fitting of experimental results in those regions where the continuum model failed to replicate them, such the central region in the necrotic core experiment, as commented in Section 3. Eventually, using averaging techniques, the phenomena incorporated at the individual level could be translated to the continuum model to have a more computationally efficient tool. We believe that this interplay between continuum and discrete models, where discrete models allow for including biological behavior rules directly into the models and they are in turn translated to continuum models for large scale simulations, is key to the successful simulation of tumor evolution across the scales. With this work we complete a first step in this framework, obtaining a validated ABM for GB evolution. In the future, the developed model could be extended to include, as commented, mechanical cell-cell and cell-substrate interactions, to evaluate their impact in the simulation outcome for replicating the experiments. Besides, there are phenomena that are better studied at the cellular scale [Gallaher et al., 2020], such as the interaction between tumor cells and the immune system, which could also be a promising field of study due to its great impact on GB progression and potential use in the design of therapies [Jain et al., 2023].

Besides, in this work we have also tackled the implementation in ABMs of boundary conditions for cells in confined regions. The usual situation in ABMs is to have a domain that is much larger than the region of interest so that the behavior at the border does not influence the quantities of interest in the study. Thus, the standard implementation of many ABMs is that cells can escape through the boundaries. To study cell evolution in microfluidic devices, we need to incorporate impermeability boundary conditions (equivalent to Neumann boundary conditions with no flux in continuum models), so we have proposed a condition for doing so in the ABM. Regarding cell escape we have included a probability of escape in the model, but it remains to relate this implementation with Robin boundary conditions used to simulate cell leakage in continuum models such as [Ayensa-Jiménez et al., 2020].

Finally, the presented methodology is limited to parameters sets for which it is feasible to determine a mathematical equivalence. In the case when this is not feasible (or highly complex), we could explore alternative methodologies to relate both models. For instance, machine learning tools, such as neural networks, could be investigated to obtain relationships between both scales and models in cases of incomplete information or noise.

## 5 Conclusions

GB is a highly invasive and aggressive brain cancer with a very poor prognosis, whose research combines all kinds of tools: *in vivo, in vitro* and *in silico*, to try to improve its treatment and provide answers to aspects that are still not understood.

In particular, *in silico* mathematical models are invaluable tools for understanding underlying mechanisms and interactions, establishing trends, and testing new hypotheses. Hence, to study the evolution of this tumor *in silico*, in this work we develop an ABM. To avoid the cumbersome calibration process we take advantage of an existent and previously calibrated continuum model for the same problem, and obtain analytical expressions that relate the parameters of both models. The resulting ABM is able to reproduce the experimental results as good as the continuum model, incorporating the inherent uncertainty of the predictions.

In summary, the main contributions of the work herein presented are:

- We have established a link between ABM and continuum models. Mathematical relations have been derived between the parameters of continuum and ABM for GB evolution This methodology avoid any calibration procedure, despite of the parameters related with experimental operation.
- An ABM for GB evolution on microfluidic devices has been developed, using the open source library Physicell. The main characteristics of GB evolution have been successfully reproduced with the proposed ABM.

However, and despite our efforts, we need more research and experimental data to understand all the complex processes involved in the evolution of the GB and its underlying mechanisms, and to develop a global model with its corresponding parameters, remains a formidable challenge. Regarding this work about the definition of ABMs for GB evolution, the long-term goal is to add other local interactions between cells (mechanical interactions for example) and individual-level phenomena that are difficult to implement in continuum models to improve the reproduction of experimental results and create complete and accurate digital twins of the *in vitro* experiments.

## Acknowledgments

The authors gratefully acknowledge the financial support from the CDTI project Chineka (EXP 00149754 / IDI-20220819). We also acknowledge the financial support from the Spanish Ministry of Science and Innovation (MICINN), the State Research Agency (AEI), and FEDER, UE through the GBM_IMMUNE project (PID2021-126051OB-C41:/AEI/10.13039/501100011033/FEDER, UE) and the FPI grant (PRE2022-102428). Finally, the authors would like to acknowledge the financial support from the Government of Aragón, Spain, through a predoctoral grant as well as the support to the research group (DGA-T62_23R).

## Appendix

### Mathematical derivations for the obtention of the parameters of the Agent-Based model

This appendix is devoted to the derivation of relationships between the functions defining an ABM of the class used in this work from those corresponding to a generic continuum model based on Reaction-Convection-Diffusion Equations (RCDEs), typically used to simulate biological processes. Note that to lighten the notation, the variables and parameters of this appendix are independent to the ones used in the main text so that the developments are self-contained.

#### Reaction-Convection-Diffusion Equation

The RCDE is a partial differential equation that describes the combined effects of reaction, convection, and diffusion in a given physical system. The general form of the RCDE for the evolution of a given field *u* = *u*(***x***, *t*) can be written as:

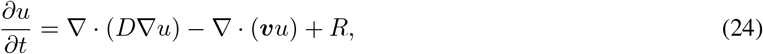

where *c* is the variable field of study, *D* the diffusion coefficient, ***v*** the drift velocity and *R* the reaction term. The reaction term, *R*, can take different forms, but in our case, we decompose it in a source and sink term, both dependent on the values of the field *u* and any other fields related with the microenvironment, *ρ*_*i*_ = *ρ*_*i*_(***x***, *t*), *i* = 1, …, *k*.

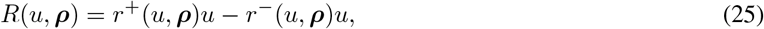

where we use the notation ***ρ*** = (*ρ*_1_, …, *ρ*_*k*_) for referring to all species. nAgent-Based model

##### A.1.1 Migration

The ABM used in this work has two parameters related to migration: the migration speed (*s*) and the migratory bias (*b*). Additionally, the velocity of directed migration has to be defined via the unit vector ***p***.

To determine the relationship between the cell migration parameters in the ABM and those of the continuum model (namely the diffusion coefficient *D* and convective velocity ***v***), we relate the parameters in both models to those of a random walk model (RWM). The derivation is based on obtaining an average velocity in both discrete models (ABM and RWM) that can be compared to the macroscopic parameters in the RCDE. First, we will develop this in the RWM and in the ABM respectively, and then we will establish the equivalences between models.

#### Random walk model (RWM)

The random walk process is defined by means of a random variable ***S***_*i*_ that is the displacement of a particle at time step *i*. Such a particle moves a distance *l* in a random direction for each time step *τ*. We set the parameter *ϵ∈* [0, 1] that determines how random or deterministic the motion of that particle is, so that the mean displacement of the particle at that time step is:

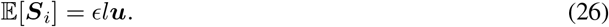

where ***u*** is a unit drift vector denoting the migration bias. Therefore, if *ϵ* = 0 the motion is totally random and if *ϵ* = 1 the motion is totally guided.

##### Proposition 1.

*Let* ***R*** *be the random vector representing the total displacement at a time point t* = *Nτ*,

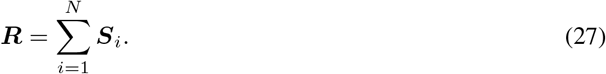

*Then, if* ***r*** =𝔼 [***R***] *is the expected value of* ***R*** *and σ*^2^ = Var(***R***) *is the variance of* ***R***, *we have*

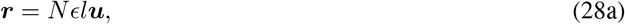

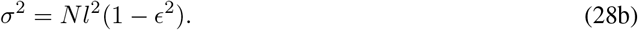

*Proof*. The expectation of the total displacement ***R*** is

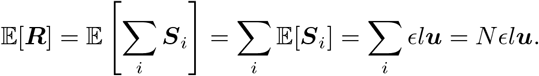

The variance of ***R*** is defined as:

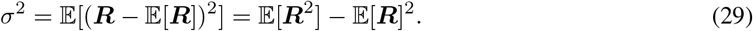

As we have already calculated the expectation of the total displacement ***R***, we develop the term 𝔼 [***R***]^2^:

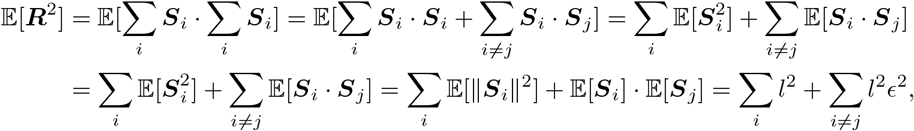

when in the last step we have used that the movements of a particle at two different time steps *i* and *j* are independent.

Finally, the expectation of the total squared displacement is as follows:

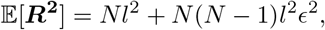

and therefore the variance of the displacement is expressed as:

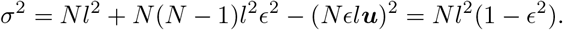

From here we can obtain the average particle velocity (which macroscopically is equivalent to the convective or drag velocity) as the expectation of the total displacement divided by the total time:

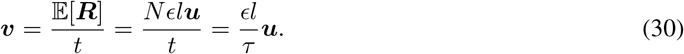

As ***u*** is a unit vector, it follows inmmediately that

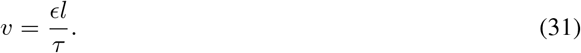

By definition (see for instance Codling et al. [2008]), the variance (*σ*^2^) is related to the diffusion coefficient *D*:

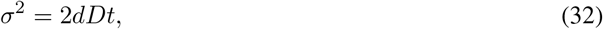

where *d* is the dimension of the space, *D* the diffusion coefficient and *t* the time. By combining equations (28b) and (32) we obtain as a final result:

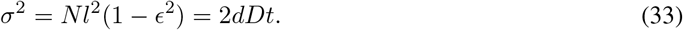

from which we obtain the relationship between the diffusion coefficient and the parameters *ϵ, l* and *τ* :

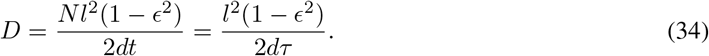

The equations (31) and (32) may be inverted to express *ϵ* and *l* in terms of *D* and *v*:

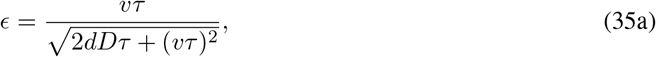

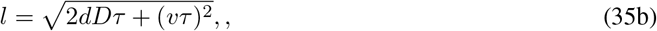

provided that *τ* is given.

##### Agent-based model (ABM)

The migration velocity in this model is given by ***v*** = *s****u***, where *s* is the modulus of the velocity and ***u*** is the vector defining the direction of cell motion according to Eq. (1):

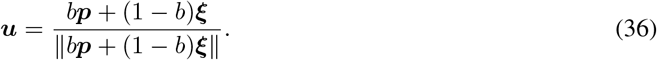

To relate the vector ***v*** to the continuum model, we consider the random vector representing the migration velocity ***V*** in terms of the random unitary vector **Ξ**

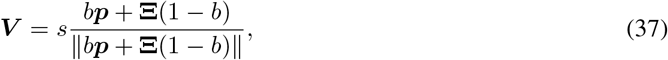

###### Proposition 2.

*Let us consider the expectation of the random vector* ***V***, ***v*** = 𝔼 [***V***], *then*

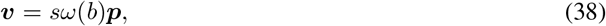

*where*

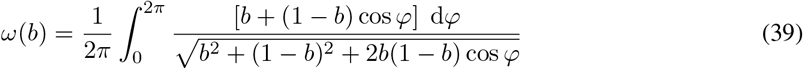

*Proof*. We first express the unit vectors ***p*** and **Ξ** using polar coordinates:

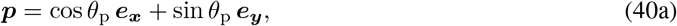

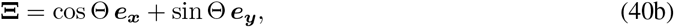

where *θ*_*p*_*∈* [0; 2*π*] is a fixed angle and Θ is a random variable with Θ*∼* 𝒰[0; 2*π*], being 𝒰 [*a*; *b*] the uniform distribution in𝒰 [*a*; *b*], that is 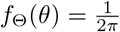.

Substituting (40a) and (40b) into equation (5), and using elementary trigonometry relations, ***V*** becomes:

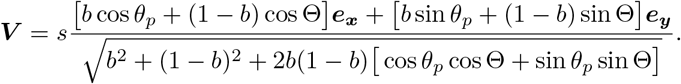

We now demonstrate that ***v****∝* ***p***, or, equivalently ***v****·* ***p***^*⊥*^ = 0. Noting that by linearity𝔼 [***V***] *·* ***p***^*⊥*^ = 𝔼 [***V****·* ***p***^*⊥*^] so that:

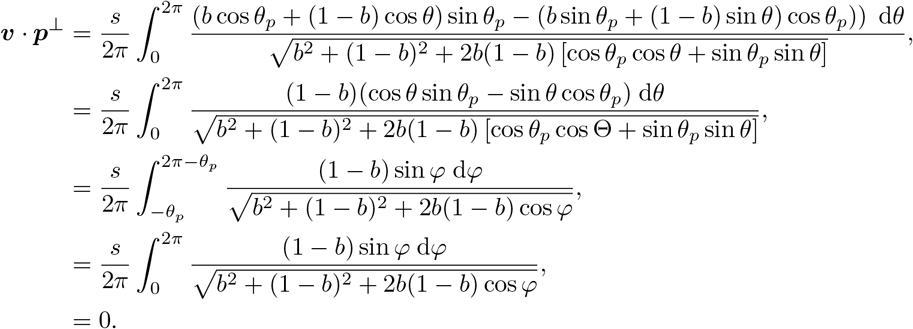

where we have used elementary trigonometric identities, the change of variable *φ* = *θ− θ*_*p*_ in the third equality and the fact that the integrand is 2*π*-periodic and odd in the two last equalities.

Now we calculate the expectation of the scalar product of ***V*** with ***d***, noting again that 𝔼 [***V***] *·* ***d*** =𝔼 [***V*** *·* ***d***] so

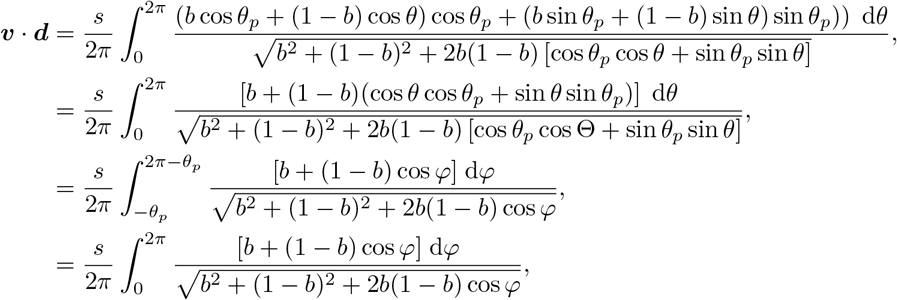

where we have used again elementary trigonometric identities, the change of variables *φ* = *θ− θ*_*p*_ at the third equality and the 2*π*-periodicity at the last one.

Hence, ***v*** = (***v*** *·* ***d***^*⊥*^)***d***^*⊥*^ + (***v*** *·* ***d***)***d*** = *sω*(*b*)***d*** with

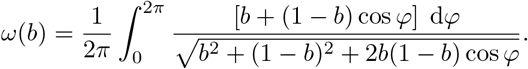

As a final result, we obtain the expected value of the migration speed vector for the agent-based model as a function of the parameter *b*, the parameter *s* and the dirft velocity ***p***

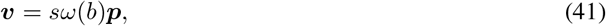

It is straightforward to see that if *b* = 0, *ω* = 0 so ***v*** = **0** and if *b* = 1, ***v*** = *s****p*** so the expected migration is in the direction of the drift vector and has modulus *s*.

##### Equivalence between models

We postulate the first-order equivalence RWM and the ABM by explicitly stating that the expected value of the migration velocity is the same for both models, that is, using Eqs. (30) and (41):

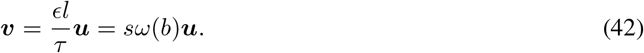

Hence, it is possible to identify both models by stating:

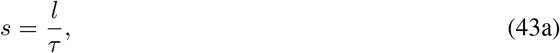

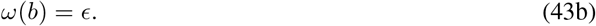

Finally, in terms of the parameters governing the RCDE. *s* and *b* are obtained substituting the expressions relating the RCDE and RWM (equations 35a, 35b) in the expressions relating the ABM and RWM (equations 43a, 43b), finally obtaining

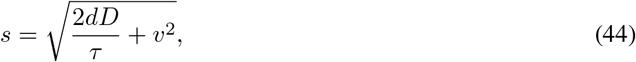

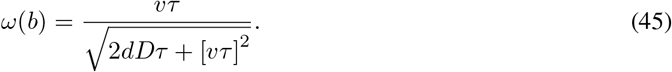

Finally, the relationship of ***p*** in the ABM to the continuum is immediate, since the migration direction is defined as:

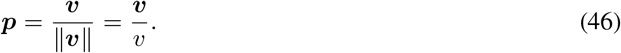

Equations (44), (45) and (46) allow to express the parameters of the discrete model in terms of the ones of the continuum one.

##### A.1.2 Proliferation and death

The ABM proliferation term can be related to the proliferation term in the reaction term of the RCDE. First, we need to obtain the cell concentration from the probabilities of transition between phases. If *n*_*t*_ is average the number of agents at a time *t* at a Reference Volume Element (RVE) *V* and the probability of agent division is *p*:

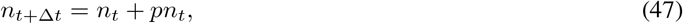

that is, the number of agents at the next time step *t* + Δ*t* equals number of agents at time *t* plus the expected number of new agents. This is formally obtained by considering that each agent replication at time *t* follows a Bernouilli distribution with parameter *p*, the number of new agents follows a Binomial distribution Δ*N*_*t*_*∼ℬ* (*n*_*t*_, *p*) and therefore E[Δ*N*_*t*_] = *pn*_*t*_. Dividing Eq. (47) by the RVE we obtain for the agent concentration at time *t, u*_*t*_:

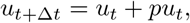

Considering that, according to Eq. (1), this probability is calculated as *p* = *P* (division) = *g*Δ*t*,

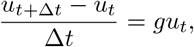

and if we take the limit when Δ*t→* 0 and we identify the cell concentration at time *t* with the solution field *u*(***x***, *t*), we obtain the continuos description of the derivative of the solution field:

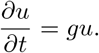

As can be seen, this equation can be identified with the proliferation term in Eq. (25) with a function dependency on the chemical species (***ρ***), so we can obtain the relationship between the parameter in the ABM and the parameters of the continuum model:

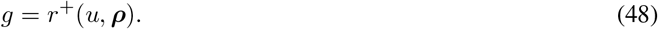

Following a procedure analogous to that of the previous section, from Eq. (2) we obtain how the concentration of dead cells changes over time in the ABM:

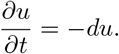

Therefore, we obtain in an analogous manner the relationship between the ABM death ratio and the reaction term associated with death in the continuum (Eq. 25):

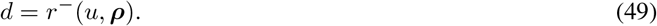

The two functions *r*^+^ and *r*^*−*^ are the growth and death rates, that depend on the microenvironment conditions, via the functions *ρ*_1_, …, *ρ*_*k*_, and on the solution field *u* itself.

##### A.1.3 Uptake and Secretion

In this section we will derive the cell consumption parameter (*U*_*ij*_) and the cell secretion parameter (*S*_*ij*_) by comparing the RCDE and Eq. (6) that we repeat here for completeness.

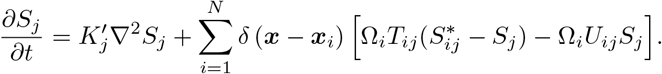

Replacing the Dirac by its discrete approximation in a certain voxel *𝒱*_*k*_, *δ*(***x*** *−* ***x***_*i*_) *≃* 1*/*Π_*k*_ if ***x***_*i*_ *∈ 𝒱*_*k*_ being Π_*k*_ = *µ*(*𝒱*_*k*_) the volume of the *k*-th voxel we obtain that.

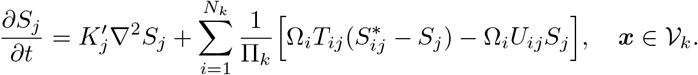

where *N*_*k*_ is the number of agents in the *k*-th voxel, Σ_k_ N_k_ = N, that is

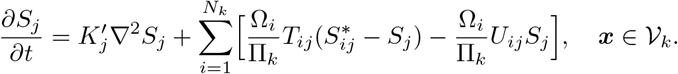

On the other hand, we have the RCDE for the evolution of each species *ρ*_*j*_, Eq. (8), is:

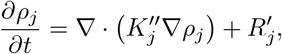

where

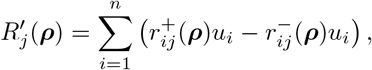

where we use the notation ***u*** = (*u*_1_, *u*_2_, …, *u*_*n*_) and ***ρ*** = (*ρ*_1_, …, *ρ*_*k*_) for referring to the population fields and the species respectively. If the diffusion process is homogeneous, it is straightforward to see that 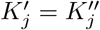 and

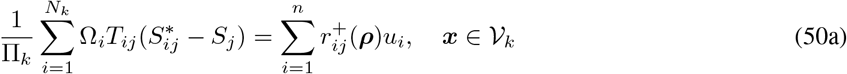

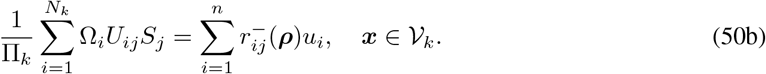

Now we consider the case where there are *n* classes of agents and all the agents belonging to one specific class have the same volume and the same behavior with respect to the species *j*. Now Eqs. (50) write

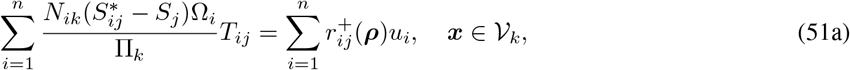

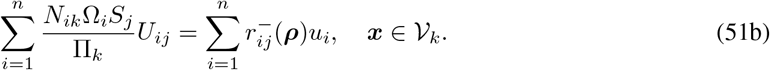

where *N*_*ik*_ is the number of agents of the *i*-th population at the *k*-th voxel. By making the identification *N*_*ik*_*/*Π_*k*_*≃ u*_*i*_, that is reasonable if Ω_*i*_*≪* Π_*k*_ and *N*_*ik*_*≫* 1, and provided that *u*_*i*_ is arbitrary, we arrive to the following identification for each population *i*:

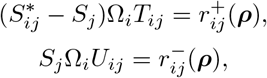

and therefore

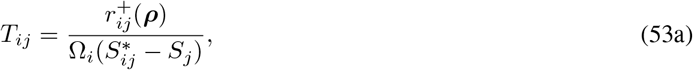

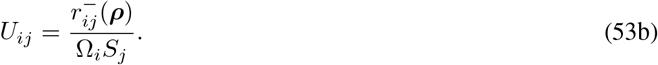

